# Engineered reciprocal chromosome translocations drive high threshold, reversible population replacement in Drosophila

**DOI:** 10.1101/088393

**Authors:** Anna B. Buchman, Tobin Ivy, John M. Marshall, Omar S. Akbari, Bruce A. Hay

## Abstract

Replacement of wild insect populations with transgene-bearing individuals unable to transmit disease or survive under specific environmental conditions provides self-perpetuating methods of disease prevention and population suppression, respectively. Gene drive mechanisms that require the gene drive element and linked cargo exceed a high threshold frequency to spread are attractive because they offer several points of control: they bring about local, but not global population replacement; and transgenes can be eliminated by reintroducing wildtypes into the population so as to drive the frequency of transgenes below the threshold required for drive. It has long been recognized that reciprocal chromosome translocations could, in principal, be used to bring about high threshold gene drive through a form of underdominance. However, translocations able to drive population replacement have not been reported, leaving it unclear if translocation-bearing strains fit enough to mediate gene drive can easily be generated. Here we use modeling to identify a range of conditions under which translocations should spread, and the equilibrium frequencies achieved, given specific introduction frequencies, fitness costs and migration rates. We also report the creation of engineered translocation-bearing strains of *Drosophila melanogaster*, generated through targeted chromosomal breakage and homologous recombination. By several measures translocation-bearing strains are fit, and drive high threshold, reversible population replacement in laboratory populations. These observations, together with the generality of the tools used to generate translocations, suggest that engineered translocations may be useful for controlled population replacement in many species.

---

Insects act as vectors for a number of important diseases of humans, animals, and plants (1). Traditional vector control is often challenging, with the degree of protection provided being proportional to the effort put into control. In addition, depending on the environment, specific methods of vector control, such as environment modification or use of insecticides, may be impractical or have undesirable side effects. A complementary strategy for disease prevention, first articulated many decades ago (2), involves using gene drive to bring about replacement of wild, disease transmitting insect populations with individuals engineered to be refractory to disease transmission, but still subject to traditional vector control (reviewed in (3-6). In a variant of this idea, population replacement has also been proposed as a method for bringing about disease prevention and/or a reduction in insect mediated damage through periodic population suppression (7, 8). This can occur when replacement results in all individuals carrying genes that cause death or failure to diapause in response to application of an otherwise benign chemical, or a seasonal change in an environmental variable such as temperature or humidity. An important appeal of these strategies is that they are species-specific and potentially self-perpetuating.

Because transgenes that mediate disease resistance or conditional lethality are unlikely to confer a fitness benefit to carriers an essential component of most population replacement strategies (see (9-12) for several non-drive based replacement strategies) is linkage with a gene drive mechanism that carries transgenes to high frequency following release. These drive mechanisms must be strong enough to spread genes to high frequency in wild populations on human timescales, while also functioning within regulatory frameworks (13-17). Central to the latter are issues of confinement and reversibility: can the spread of transgenes to high frequency be limited to locations in which their presence is sought, and can the population be restored to the pre-transgenic state?

An important characteristic of any gene drive mechanism that relates to the above questions is its level of invasiveness: its ability to increase in frequency both at the release site and in surrounding areas linked to the release site by various levels of migration, when introduced at various population frequencies. Low threshold gene drive mechanisms require that only a small fraction of individuals in the population carry the drive element in order for spread to occur locally (18, 19). Examples include engineered *Medea* chromosomal elements (20-22), several other possible single locus chromosomal elements (23), site-specific nucleases that home into their target site (24-29), and site-specific nucleases that result in sex ratio distortion (30). These mechanisms are predicted to be invasive because low levels of migration of drive element-bearing individuals into areas outside the release area may, depending on the threshold and the migration rate (18, 19, 31), result in these areas being seeded with enough transgene-bearing individuals for drive to occur. Low threshold, invasive gene drive mechanisms are attractive when the goal is to spread transgenes over a large area, and migration rates between the release site and surrounding areas of interest are low. However, for these same reasons, it is likely to be challenging to restore the population to the pre-transgenic state if desired. Given the intense scrutiny with which releases of insects engineered to suppress population numbers while ultimately disappearing from the population have been greeted (15-17), gene drive mechanisms that have a limited capacity to spread, and that can easily be eliminated from the population, thereby restoring the population to a pre-transgenic state, may be useful in some contexts.

High (or higher) threshold gene drive mechanisms require, as their name implies, that transgenes make up a much larger fraction of the total insect population (important examples range from 15-70%) before gene drive occurs. Below this frequency transgenes are instead actively eliminated from the population. In short, these drive mechanisms behave as a frequency-dependent bistable switch. High transgene frequencies are needed to initiate drive at the release site, limiting the possibility that unintended release of a few individuals could initiate replacement. Once replacement has occurred at the release site, spread to high frequency in areas connected to the release site by low levels of migration is prevented because the transgene never reaches the threshold frequency needed for drive. Finally, transgenes can be eliminated from the population if the release of wildtypes results in the frequency of transgenics being driven below the threshold required for drive.

A number of gene drive mechanisms that could in principal bring about local and reversible population replacement have been proposed. Examples include a number of single locus gene drive mechanisms (23, 32, 33), reciprocal chromosome translocations, inversions and compound chromosomes (34), and several forms of engineered underdominance (23, 35-39) (40). One of these, UD^MEL^ (double *Medea*), has recently been shown to drive reversible population replacement into populations of wildtype Drosophila (38). A second system has been shown to drive high threshold population replacement in Drosophila in a split configuration (40). In each of these systems gene drive occurs when transgene-bearing chromosomes experience frequency-dependent changes in fitness with respect to non-transgene-bearing counterparts, with the former having high fitness at high frequency and lower fitness at low frequency. These systems all rely, in one way or another, on the phenomena of underdominance, in which transgene-bearing heterozygotes (or some fraction of them or their progeny) have a lower fitness than either homozygous wildtypes or homozygous transgenics (or transgene-bearing trans-heterozygote in some three allele cases). If the frequency of one allele or pair of alleles or chromosome type is above a critical threshold it spreads to genotype, and in some cases allele fixation. Conversely, if it falls below the critical threshold it is lost in favor of the other allele or chromosome type, usually wildtype. In broad outline, this behavior occurs because when transgene-bearing individuals are common they mate mostly with each other, producing transgene-bearing offspring of high fitness (high survival and/or fecundity), while wildtypes mate mostly with transgene-bearing individuals, producing a preponderance of heterozygous offspring of low fitness (inviable and/or with reduced fecundity). However, when the frequency of wildtypes is high the tables are turned, with transgene-bearing individuals producing high frequencies of unfit heterozygous progeny, and wildtypes producing a high frequency of fit homozygous progeny.

Here we focus on the use of engineered reciprocal chromosome translocations as a high threshold gene drive mechanism. Reciprocal chromosome translocations were the first gene drive mechanism proposed (2). Their structure and genetic behavior are illustrated in Figure 1A. A reciprocal chromosome translocation results in the mutual exchange of DNA between two non-homologous chromosomes (41). Provided that the translocation breakpoints do not alter the expression and/or function of nearby genes, translocation heterozygotes and homozygotes can in principal be phenotypically normal. Thus, phenotypically normal, naturally occurring translocation-bearing individuals are found in populations of many species (42), including humans (43, 44). However, translocation heterozygotes are usually semisterile, producing a high frequency of inviable offspring. This occurs because meiosis in a translocation heterozygote can generate a variety of different products. Three patterns of segregation are possible: alternate, adjacent-1 and adjacent-2 (Figure 1A). While alternate segregation leads to the production of gametes with a full genome complement, adjacent-1 and adjacent-2 segregation lead to the production of aneuploid gametes, resulting in the death of progeny that inherit an unbalanced chromosome set. In many species alternate and adjacent-1 segregation occur roughly equally, with adjacent-2 segregation being rare (45, 46). In such species progeny genotypes and survival phenotypes resulting from crosses between translocation-bearing individuals and wildtypes are as illustrated in the Punnett square in Figure 1B. Progeny with unbalanced genotypes die, while balanced translocation heterozygotes, translocation homozygotes, and homozygous wildtypes survive.

**Figure 1.**
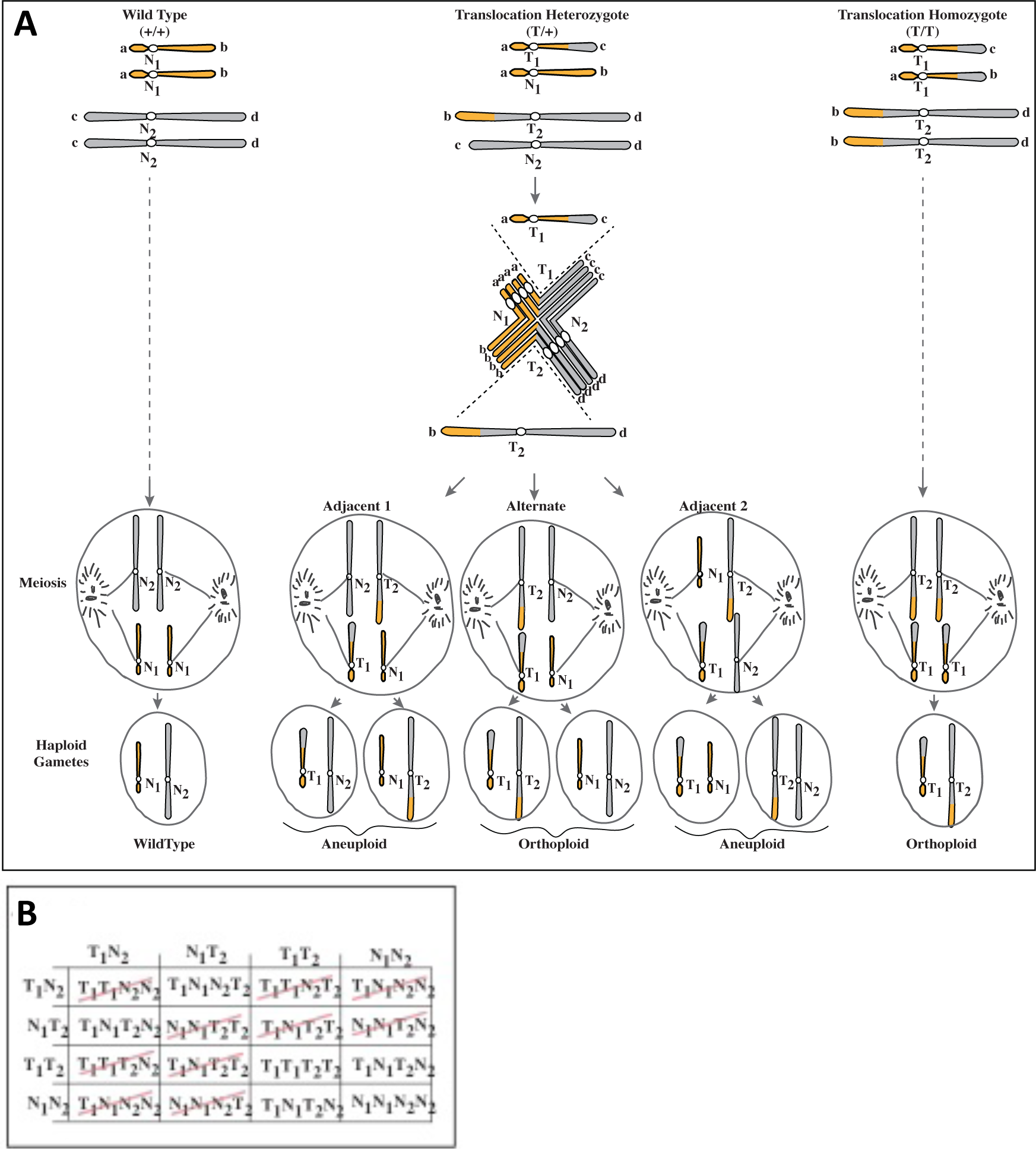
Gamete and zygote genotypes associated with the presence of a reciprocal translocation. Wildtype chromosomes N1 and N2, and translocation chromosomes T1 and T2, are indicated. (A) One chromosome type (a) is indicated in yellow. A second chromosome type (b) is in gray. Gamete types generated by wildtype (+/+), translocation heterozygotes (T/+), and translocation homozygotes (T/T) are indicated. (B) Gamete and zygote genotypes possible in crosses involving a translocation are indicated. Inviable genotypes are indicated by a red line.

In 1940 Serebrovski proposed that the release of homozygous translocation-bearing males could be used to drive population suppression because many progeny would be semisterile, thereby driving down population fitness over multiple generations (47). He, Dobzhansky (48), and later Curtis (2), also noted that the frequency of translocations lacks a stable internal equilibrium, with either wildtype or translocation-bearing chromosomes spreading to fixation in an isolated population through natural selection (differential survival of the relevant chromosome type) if their frequency rose above 50%, for a translocation with no fitness cost to carriers. Curtis proposed that if a gene beneficial to humans could be linked to the translocation breakpoint, this behavior of translocations could be used to spread the gene into the wild population. Whitten subsequently noted that the same approach could be used to spread a trait conferring conditional lethality, which could be used to bring about population suppression (7). More recent modeling work has highlighted the potential of translocations for bringing about local, but not global population replacement, and the ease of reversal (19).

Though it is clear from evolutionary studies that translocations can become fixed in populations (42), efforts to directly bring about population replacement using translocations created in the lab have not been successful (34, 49-51). There may be several reasons for this. First, translocation-bearing individuals (particularly homozygotes) generated in the past typically had very low fitness, probably at least in part because they were generated using X-rays, which can result in a high frequency of background mutations. Second, more recently it has become clear that chromosome positioning and structure in the nucleus can play a role in determining large-scale patterns of gene expression, and that chromosome translocation can result in changes in the patterns of gene expression (52, 53). These latter observations leave it fundamentally unclear whether translocation-bearing individuals of high fitness can be easily generated, even if the breakpoints involved are located in gene deserts. For example, it could be that phenotypically normal translocation-bearing individuals observed in nature simply represent the relatively rare cases in which chromosome rearrangement does not result in fitness being compromised. To explore these issues, and to determine if translocation-based gene drive can be used to bring about population replacement, we first use modeling to explore the relationship between variables such as introduction frequency, fitness cost, and reciprocal migration with non-target populations containing widltypes, for the ability of a translocation to spread, and the equilibrium frequencies achieved in replaced and surrounding populations. We then describe a general approach to generation and identification of site-specific reciprocal chromosomal translocations. Finally, we provide the first demonstration that engineered translocations are capable of bringing about threshold-dependent population replacement, in *Drosophila melanogaster*.

## Some predicted characteristics of translocation-based gene drive

Early modeling work by Serebrovskii and Curtis showed that if a translocation results in no fitness cost to carriers, and is present in a population experiencing no incoming migration of wildtypes, it will spread to allele fixation when present at population frequencies greater than 50%, and will be eliminated when present at lower frequencies (2). Curtis also noted briefly that translocations that resulted in a fitness cost to carriers could still spread to allele fixation, but the threshold introduction frequency would be increased (2). Given the past failures to bring about translocation-mediated population replacement noted above, and the likelihood that chromosome translocation itself and/or the GOI placed at the breakpoints will result in some fitness cost to carriers, we sought to understand more generally how fitness cost affects translocation spread. The time to allele fixation is particularly relevant for contexts in which the goal is to ultimately bring about population suppression in response to a seasonal variable such as temperature or humidity.

In figure 2A we illustrate the relationship between fitness cost, introduction frequency and time to translocation allele fixation (approximated as the point at which >99% of individuals carry at least one translocation copy), for a single introduction into an isolated population. The plot illustrates several important points. First, whenever translocations spread, they spread to fixation relatively quickly, with the time needed being inversely related to the introduction frequency. Second, translocations that confer large fitness costs to carriers can also spread rapidly, so long as the introduction frequency is increased. The plot in Figure 2B illustrates a related case in which the translocation is introduced over three generations at the specified frequency. It shows that with modest extra effort rapid drive can be achieved, even for very high fitness costs. While these introduction frequencies represent a large percentage of the wild population, they are still much lower than those used in self-limiting genetic population suppression strategies such as SIT and RIDL (54), and unlike SIT and RIDL, result in sustained changes to the population.

**Figure 2.**
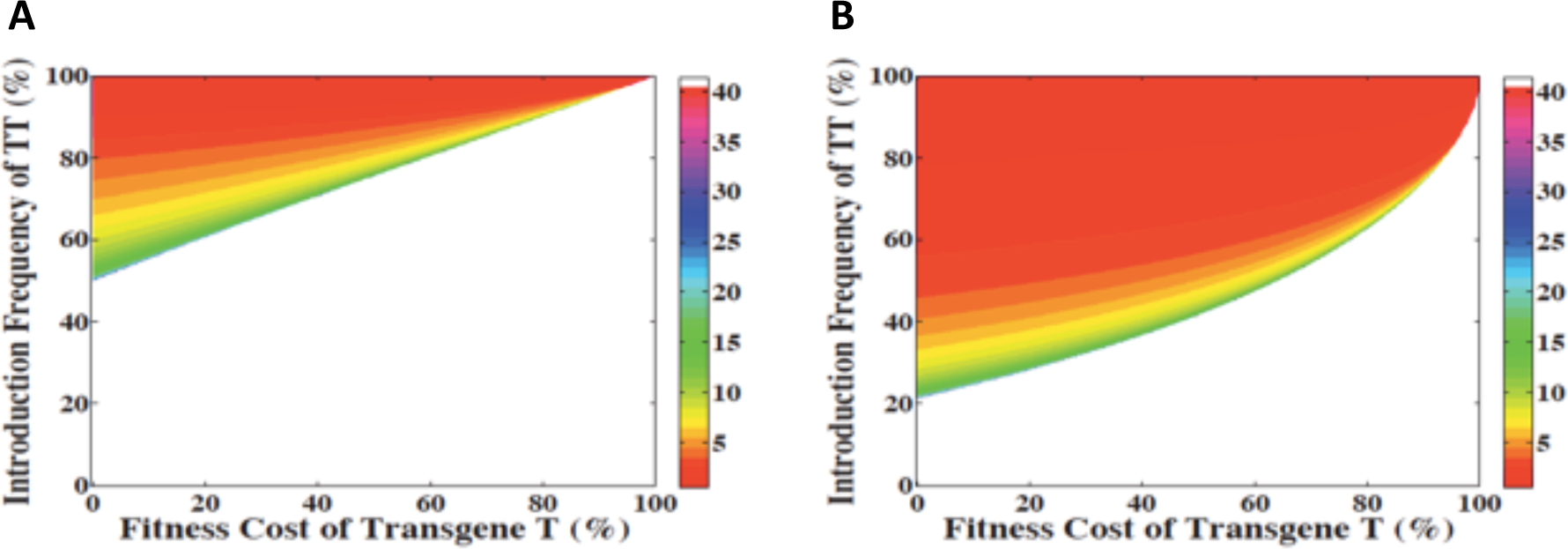
Engineered reciprocal translocations are predicted to show threshold-dependent gene drive and bring about local population replacement. A discrete generation, deterministic population frequency model of translocation spread through a single population for varying introduction frequencies and fitness costs for one (A) or three (B) introductions at the specified frequency. The heatmap indicates the number of generations required for the translocation to reach fixation (i.e., >99% of the total population) for all combinations of fitness cost and introduction frequency.

In real world scenarios other than initial field-testing ‐ in which population isolation will be essential ‐ there is likely to be some level of reciprocal migration between the target area (source population 1) and surrounding areas (population 2) containing wildtypes. Marshall and Hay showed that for realistic population sizes (>1000 individuals), there are no reciprocal migration rates that support population replacement in a second, wildtype-containing population (population 2) linked to a source population (population 1) in which replacement is initiated. Due to the high frequency of death among the progeny of translocation-bearing individuals that mate with wildtype, the frequency of translocation-bearing individuals in population 2 never rises to a level that supports drive (see also Figure 3A, C). Instead, when migration rates are high (~6.8%, or lower when the translocation is associated with a fitness cost), translocations are eliminated from both populations (19). Here we consider a related question: what effect does reciprocal migration have on the characteristics of population replacement in the target population, and the genotypic composition of neighboring populations linked by migration, in which drive does not occur?

**Figure 3.**
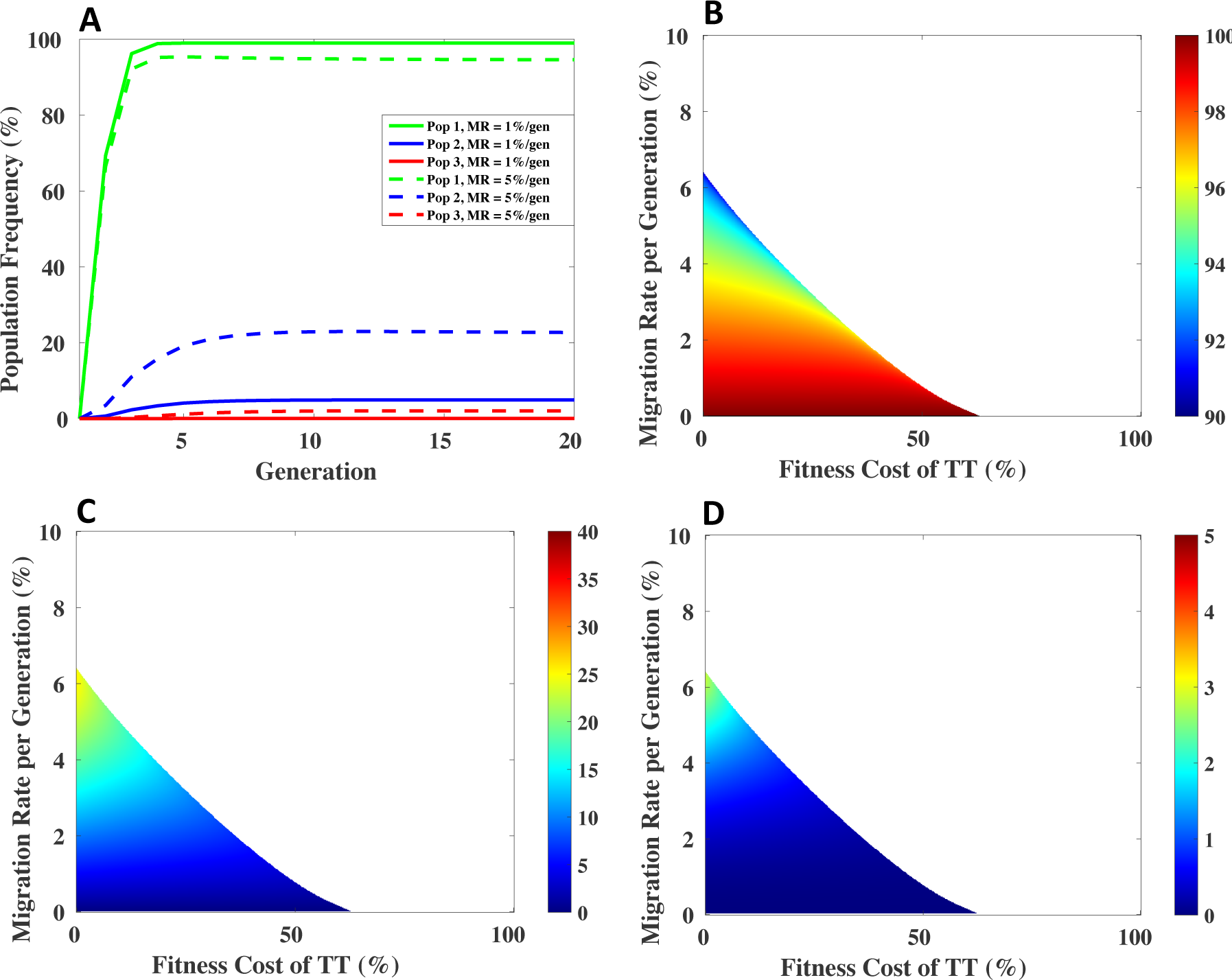
Translocation dynamics in a linear, three population migration model. (A) Population frequency of a translocation with no fitness cost, introduced into population 1 using three consecutive releases of translocation-bearing homozygotes. Populations 1-3 are linked through a linear chain of migration of 1% (solid lines) or 5% (dashed lines). (B-D) Equilibrium frequency of translocation bearing individuals over a range of fitness costs and migration rates for each of the three linked populations 1 (B), 2 (C), and 3 (D), respectively. For all three populations increasing fitness cost has little effect on the equilibrium frequency at low migration rate and increased effects at higher migration rates. In contrast, migration rate has a much stronger effect on equilibrium frequency independent of fitness cost as seen by the color gradient shifts. Note that the equilibrium frequency varies between 90-100%, 0-25%, and 0-3% in the target population (population 1), population 2, and population 3, respectively.

We consider a specific scenario in which three populations are linked in series: the target population (population 1) is linked to a second population consisting initially of wildtypes (population 2) through migration; population 2 is also linked through migration to a third population consisting initially of wildtypes (population 3), which is not linked directly with population 1. We ask what the equilibrium frequencies are in each population for different levels of migration? In the case of a low threshold gene drive mechanism such as *Medea* or homing by a HEG, the equilibrium frequency in population 1 will approach fixation since these drive elements spread invasively into surrounding populations connected to the target population by low levels of migration. In contrast, the situation for high threshold gene drive mechanisms is fundamentally different since wildtypes will, by definition, always be present in surrounding non-target populations in which transgene levels sufficient for drive are not achieved. Previous modeling studies of underdominant systems have noted that the presence of reciprocal migration can result in internal equilibria containing both wldtype and underdominant alleles (36, 37) (55). Here we consider the case of reciprocal translocations specifically.

Figure 3A illustrates a specific scenario, in which a translocation with no fitness cost is introduced into population 1 at a frequency of 70%, and is connected to a similarly sized population 2 by a migration rate of 1%. Population 2 is connected to a similarly sized population 3 by the same migration rate. The translocation spreads to high frequency (99%) in population 1, but not to allele or genotype fixation, since wildtypes are introduced into population 1 each generation. Translocation-bearing genotypes are also present at modest levels (<5% (4.954%) in population 2, and <1% (0.08116%) in population 3. Figure 3A also illustrates an identical scenario in which the migration rate is now 5%. In this case the translocation equilibrium frequency is <95% (94.55%) in population 1, <23% (22.58%) in population 2, and ~2% (2.031%) for population 3. The general relationship between fitness cost, migration rate and equilibrium frequency in population 1 is illustrated in Figure 3B. The highest level of incoming wildtype migration that can be tolerated for a translocation with no fitness cost (~6.8% / generation) results in an equilibrium translocation genotype frequency of ~90% in population 1. Decreased levels of migration result in correspondingly higher equilibrium frequencies, which approach fixation as the migration rate falls to zero (as in Figure 2). Populations 2 (Figure 3C) and 3 (Figure 3D) show the opposite behavior. As migration rate increases, the fraction of translocation-bearing individuals increases in population 2, reaching a maximum of ~25% for a translocation with no fitness cost and migration rate of 6.8%. However, for similar migration rates the fraction of translocation-bearing individuals in population 3 is dramatically reduced. Increased fitness costs result in a minimal decrease in equilibrium translocation frequency in all three populations (Figure 3B-D).

These observations illustrate a fundamental set of tradeoffs associated with high threshold gene drive. While drive can be spatially limited to a single population, this comes with a cost: the continuous introduction of wildtypes from neighboring populations, which keeps the equilibrium frequency of transgene-bearing individuals below 100%. Depending on the disease system being considered, the presence of some level of non-transgene-bearing individuals within the target area may have important epidemiological consequences, as a residual population of wildtype mosquitoes may be capable of sustaining transmission, although this remains to be investigated. Population suppression following activation of condition-dependent lethality may also be challenging in the face of significant levels of wildtype migration. Finally, the presence of some level of translocation-bearing individuals outside the target area may have regulatory implications even if these levels are insufficient for drive. That said, any such issues are likely to be local since the decrease in frequency of drive element-bearing individuals in underdominant systems drops off rapidly in a series of linked populations (Figure 3B-D). Together, these observations suggest that high threshold gene drive is likely to be most epidemiologically effective and able to satisfy regulatory requirements relating to the presence and movement of transgene-bearing organisms within target areas circumscribed by significant barriers to migration.

## Engineering Reciprocal Translocations in Drosophila

Cells or organisms carrying translocations with defined breakpoints have recently been generated using several strategies. One set of approaches begins with two non-homologous chromosomes that each have a different transgene-bearing cassette inserted at a specific position. Recombination between the two chromosomes to generate a translocation is then driven by FLP/FRT recombination (56), Cre/*loxP* recombination (57, 58), or homologous recombination following double-stranded break creation within the transgene cassettes using a site-specific nuclease (58-60). Translocations have also been generated in completely wildtype backgrounds, following Crispr/Cas9-mediated cleavage of two otherwise wildtype chromosomes followed by non-homologous end joining (61-63). In this latter case, PCR-based methods were used to identify pools of cells or individuals carrying translocations.

We sought to create translocations using a variant of the approach described by Egli et al. in which homologous recombination between two chromosomes follows double-stranded break creation using the rare-cutting site-specific nuclease I-SceI (58). However, rather than use their approach for identification of potential translocation bearing individuals, which involves scoring for the loss of the marker *y*^+^ in an otherwise a *y*^−^ background, we created a system in which recombination results in the creation of a dominant marker. This approach can be used in otherwise wildtype genetic backgrounds, in diverse species.

Two constructs (A and B) were generated (Figure 4B). Each construct includes several components. These include (from left to right) a transformation marker (the *white* gene); a location that could be used as an insertion point of a gene of interest (GOI); a promoter that drives the expression of a dominant florescent marker, either ubiquitously (the Opie2 viral promoter, (64) or in oenocytes (65); a splice donor site, and two stretches of DNA used as substrates for homologous recombination, annotated as UVW and XYZ, each roughly 670bp in length. These DNA fragments were derived from the mouse IgG locus, and thus lack homology with the Drosophila genome. Two target sites for the rare cutting homing endonuclease I-SceI were inserted between UVW and XYZ. To the right of these elements were positioned a splice acceptor, a promoterless reporter gene (GFP or dsRed), and a phiC31 recombination attB site.

**Figure 4.**
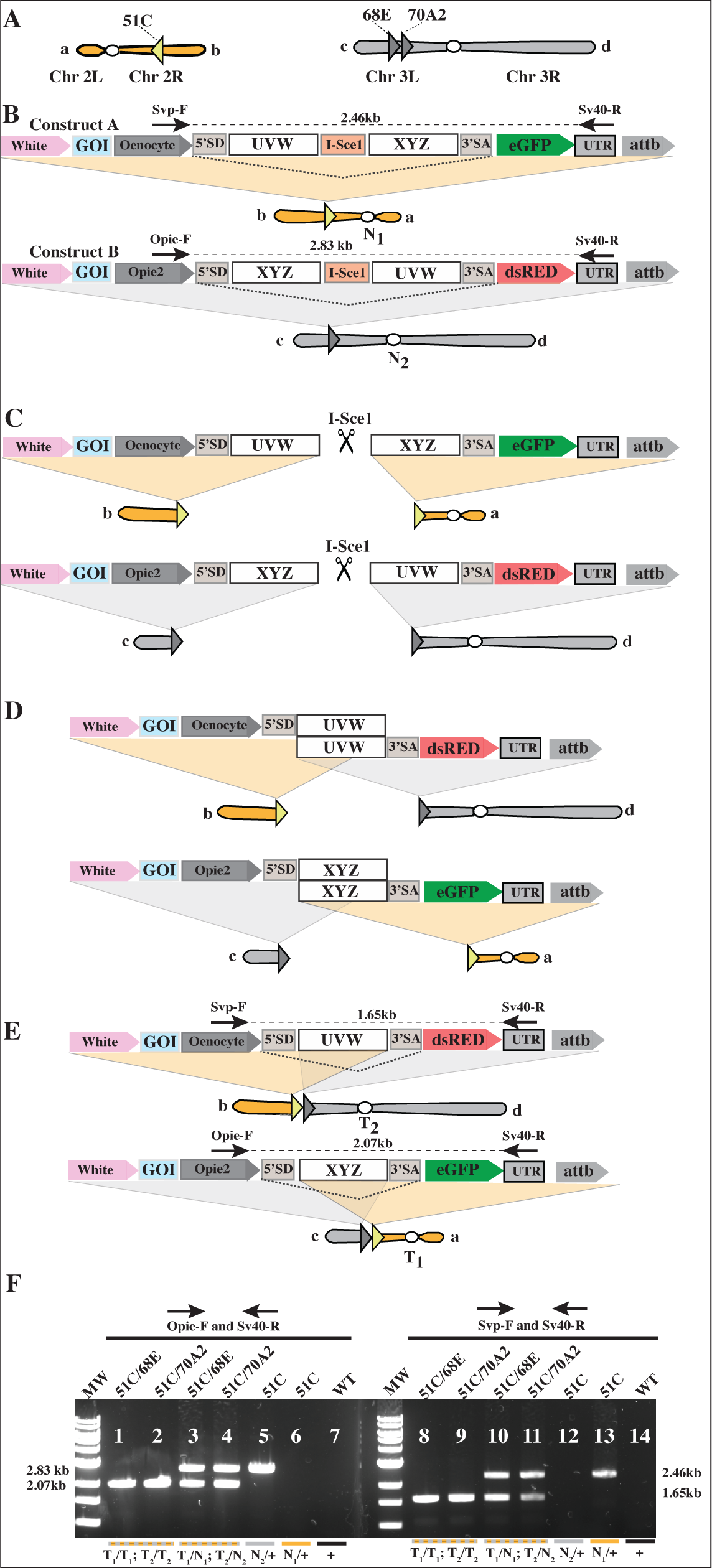
Generation of reciprocal translocations in Drosophila. (A) Approximate location of the attP sites used for transgene insertion; orientation with respect to the centromere are indicated by triangles. (B) Components of each starting transgene cassette. Construct A is inserted on the second chromosome and construct B on the third chromosome. Components are as indicated in the text. (C) I-Sce-dependent cleavage results in a double-stranded break in each transgene-bearing chromosome. (D) Alignment of broken chromosome ends occurs using homologous sequences UVW and XYZ. (E) Recombinant chromosomes are generated by homologous recombination using sequences UVW and XYZ. (F) Agarose gel image is shown of PCR amplification products generated from different genotypes: translocation homozygotes (T1/T1; T2T2); translocation heterozygotes (T1N1; T2N); individuals carrying only the 51C starting chromosome insertion (N1/+); or the 68E and 70A2 starting chromosome insertion (N2/+). Primers used, and expected amplification product sizes, are indicated in B and E.

These constructs were introduced into flies at three separate attP locations: construct A at 51C on chromosome 2, and construct B at 68E or 70A2 on chromosome 3 (Figure 4A). The attP insertion sites at 51C and 68E lie some distance from annotated genes, while the 70A2 site lies within a cluster of tRNA loci. Both constructs were oriented in the same direction with respect to their centromeres (Figure 4A). The constructs were designed so that flies bearing construct A, located on the second chromosome, would express the svp-driven eGFP marker, while construct B, located on the third chromosome, would express the opiap2-driven dsRED marker (Figure 4B). Transgenics for construct B behaved as expected, and were dsRED positive throughout their body. However, transgenics for construct A had no detectable GFP expression. The basis for this is unclear, but could be due to inappropriate splicing of the XYZ-UVW sequence in this construct. Regardless, as illustrated below, one marker is sufficient to identify translocation-bearing individuals.

To generate translocation-bearing individuals we created stocks doubly homozygous for constructs A and B (51C; 71A2 or 51C; 68E). These were then mated with flies that express I-SceI under the control of the Hsp70 heat shock promoter (66). Progeny carrying all three transgenes were subjected to multiple rounds of heat shock during larval stages and as adults. Adults were outcrossed to wildtype, and progeny examined under a fluorescent dissecting scope. In a number of individuals strong ubiquitous GFP expression was observed. This is the predicted outcome if I-SceI expression results in cleavage of both transgene-bearing chromosomes (Fig. 4C), followed by homologous recombination between XYZ-and UVW-bearing ends of the two different chromosomes (Fig. 4D,E). Putative translocation heterozygotes (T_1_/+; T_2_/+) were individually mated to wild type individuals (+/+; +/+) to generate males and female translocation heterozygotes (identified as GFP-expressing). These were mated with each other to generate putative translocation homozygotes (T_1_/ T_1_; T_2_/ T_2_). PCR and sequencing of products from genomic DNA of these individuals was used to demonstrate that these individuals were homozygous for both translocation products (Methods and Figure 4F).

To explore the genetic behavior of translocation-bearing chromosomes and the fitness of carriers we carried out a number of crosses and quantified progeny genotype (Table 1). Stocks consisting of translocation homozygotes appeared generally healthy as adults, and survival from egg to adult was 96% of that observed for the Canton S (CS) wildtype stock. In contrast, crosses between males or females heterozygous for the translocation and wildtype resulted in semisterility, with only about 50% of progeny surviving to adulthood, and 50% of the survivors being translocation heterozygotes. These are the expected results if alternate and adjacent-1 segregation occur with equal frequency in translocation-bearing individuals during meiosis, resulting in the production of 50% aneuploid gametes (Figure 1B). Finally, for each translocation type we also carried out crosses between male and female translocation heterozygotes. Only 37.5% of progeny are predicted to survive, due to the large fraction of zygotes carrying unbalanced chromosome complements. However, many of the survivors (83%) are predicted to carry one or two copies of the translocation (Figure 1B). The levels of embryo survival and percentage of adults carrying the translocation were in good agreement with these predictions (Table 1). Together, these observations suggest that the translocation-bearing strains are fit (notwithstanding the expected semisterility), at least to a first approximation. These points notwithstanding, fitness measurements such as these are not sufficient to know that frequency-dependent drive will occur. This is well illustrated by the results of Curtis and Robinson, who found that a 2;3 translocation strain generated with X-rays, which had homozygous viability and fertility equivalent to wildtype in crosses such as those described above, was unable to drive population replacement, even when introduced at a 9:1 translocation:wildtype ratio (49).

**Table 1.**
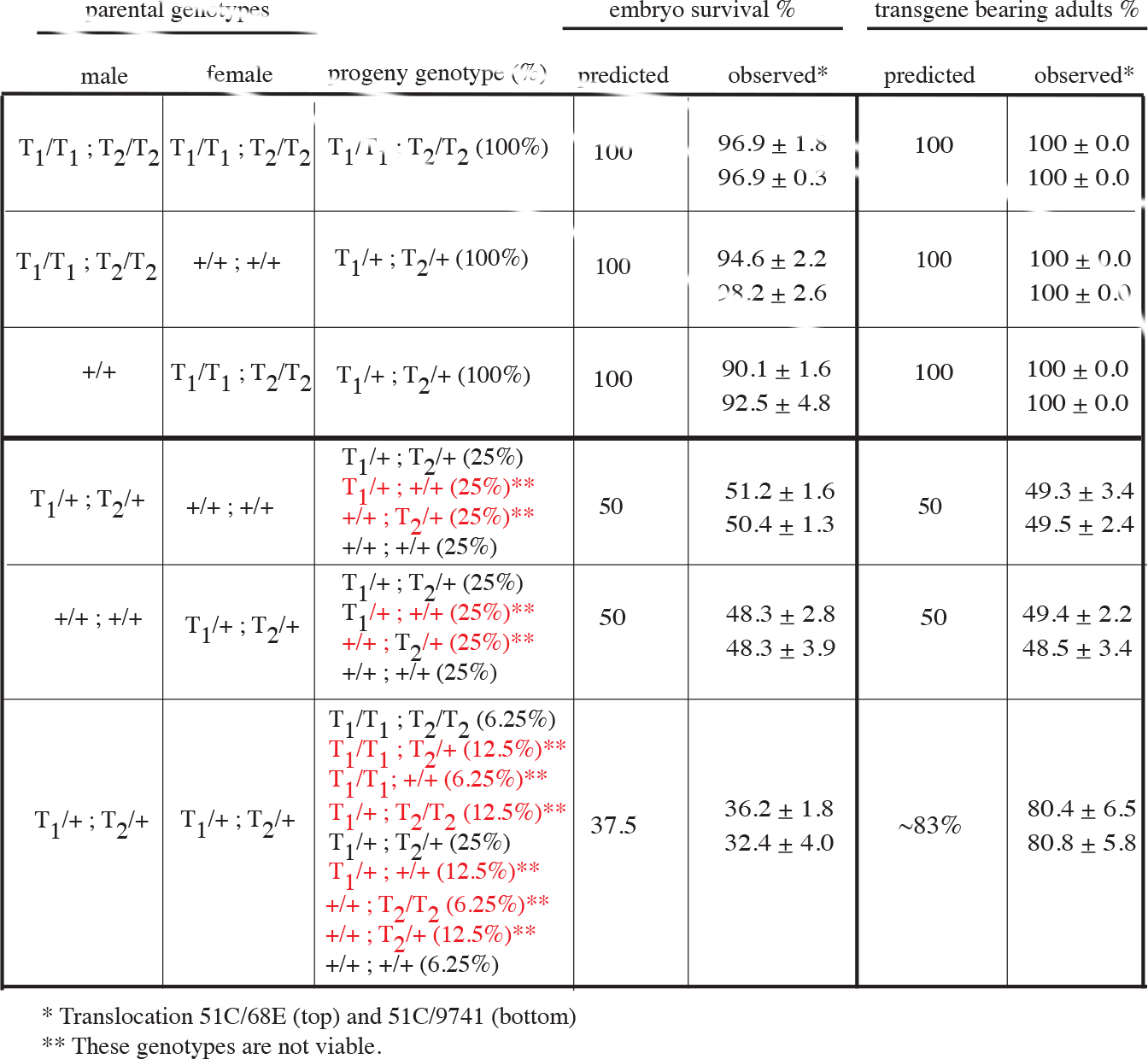
Behavior of translocations in crosses to various genotypes. Crosses between parents of specific genotypes ‐ wild-type (+/+; +/+), translocation heterozygotes (T_1_/+; T_2_/+), and translocation homozygotes (T_1_/T_1_; T_2_/T_2_), were carried out. Embryo survival (fifth column from right) and percentage of translocation-bearing adults (rightmost column) were independently quantified. The top number in each column shows results for the 51C/68E translocation; the bottom number shows the results for the 51C/70A2 translocation. ** Indicates unviable genotypes. Embryo survival was normalized with respect to percent survival (± SD) observed in the *w^1118^* stock used for transgenesis (methods).

For population replacement experiments we first introgressed our translocation-bearing systems, 51C; 70A2 and 51C; 68E flies, with Canton S (CS) for 8 generations, so as to minimize background genetic differences between translocation-bearing and wildtype strains. Translocation-bearing individuals were then backcrossed to each other to create homozygous stocks. We initiated population cage experiments by introducing translocation-bearing males and virgin females into cages along with Canton S males and virgin females of similar age. A number of different introduction frequencies were tested, in triplicate. These included frequencies predicted to be super-threshold (80%, 70%, 60%), and sub-threshold (20%, 30%, 40%). Populations were then followed for 14 generations, with the frequency of translocation-bearing individuals noted each generation.

Results of these experiments are summarized in Figure 5A,B (solid lines). For both translocation-bearing strains, all nine releases at frequencies lower than 50% resulted in elimination of the translocation from the population. Conversely, introductions at frequencies greater than 50% resulted in translocation-bearing genotypes spreading to high frequency. These results are generally consistent with the modeling predictions. However, the dynamics of drive are clearly distinct from those predicted for translocations that lack a fitness cost (dotted lines in Figure 5A,B). When translocations were introduced at predicted super-threshold frequencies spread was slower than expected for a translocation with no fitness cost. Sub-threshold releases also resulted in lower initial translocation frequencies than expected, and this was generally followed in later generations by a modestly decreased time to elimination as compared with a translocation with no fitness cost (except at the 20% introduction frequency).

**Figure 5.**
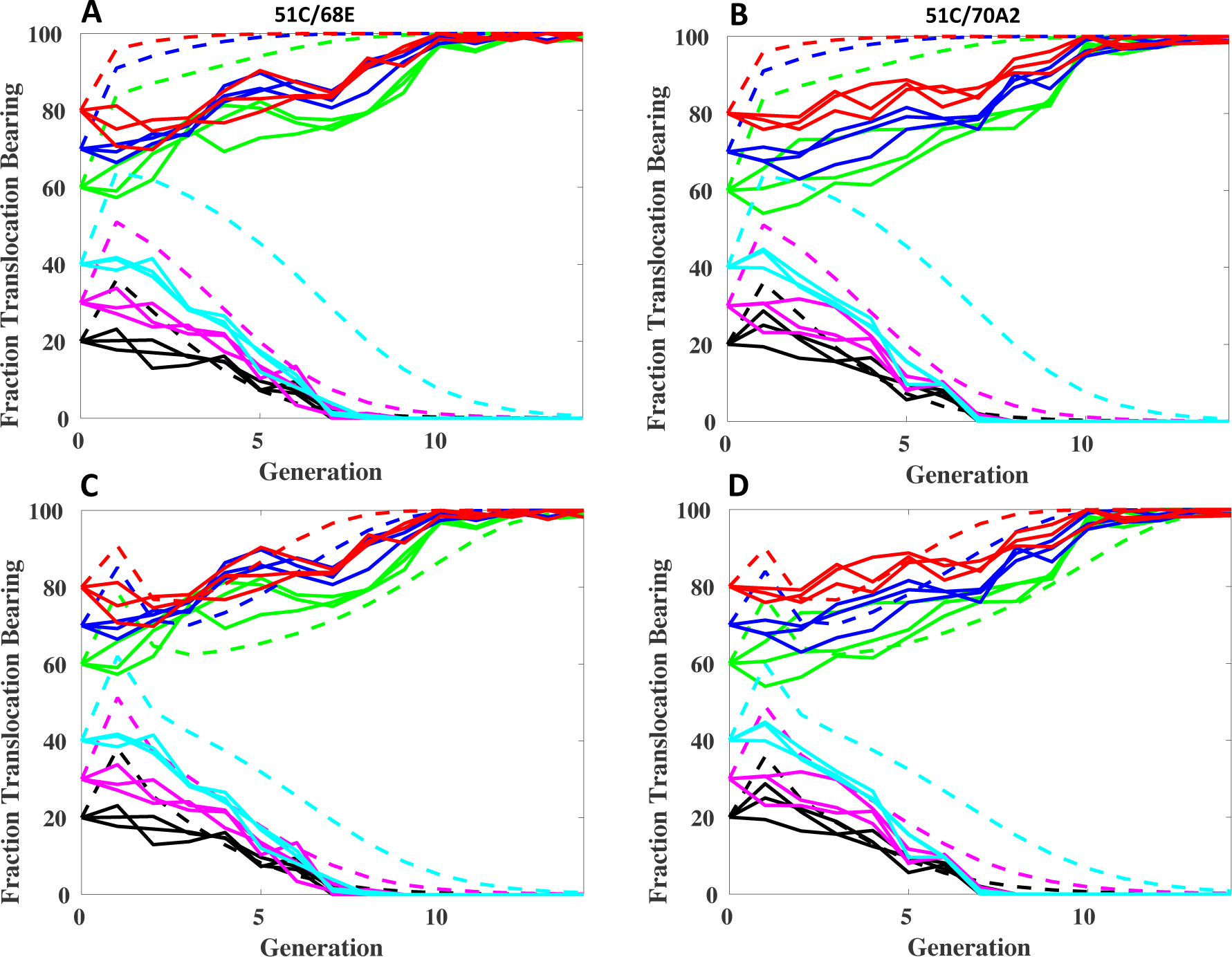
Dynamics of translocation-based population replacement, and predictions from zero fitness cost, and best fit models. (A, B) Population frequency of the adult population having the indicated translocation is plotted versus generation number for a number of homozygous translocation release ratios: 80%, 70%, 60%, 40%, 30% and 20%. Solid lines indicate observed population frequencies, and dashed lines indicate predicted translocation-bearing genotype frequencies for an element with no fitness cost. (C, D). The same data as in (A, B) but plotted along with dynamics predicted based on a best fit model described in the methods and text.

To understand these dynamics, we fitted the experimental data with our previously described deterministic model framework (19) using a range of different fitness cost models (Methods). By comparing the Akaike Information Criterion (AIC) values for each of these fitness cost models we found the best fitting model for the observed population dynamics to be one in which the relative fitness of homozygotes having the translocation is time-dependent, with the relative fitness of these individuals rapidly increasing over time, at first rapidly and converging upon some higher value as described by an exponential function. Calculations of fitness parameters for translocation system 1 suggest an initial relative fitness of transgenic homozygotes of 0.0004 (95% CrI: 0-0.0019) relative to wild-types in generation 1 (the first progeny generation post adult introduction), rising to a relative fitness of 1.51 (95% CrI: 1.48-1.53) in subsequent generations. Calculations suggest an initial relative fitness of transgenic heterozygotes of 1.23 (95% CrI: 1.14-1.31) relative to wild-types, falling slightly to a relative fitness of 1.05 (95% CrI: 1.02-1.08). Calculations for translocation system 2 suggest an initial relative fitness of transgenic homozygotes of 0.0003 (95% CrI: 0-0.0016) relative to wild-types, rising to a relative fitness of 1.52 (95% CrI: 1.50-1.55) in subsequent generations, and an initial relative fitness transgenic heterozygotes that remains fairly constant: 1.12 (95% CrI: 1.05-1.18) at the beginning of the experiment and 1.11 (95% CrI: 1.08-1.14) at the end of the experiment.

While speculative, the initial very low fitness of homozygotes in generation 1 could reflect the fact that these individuals must derive from homozygous translocation parents. Our analysis of fitness presented in table 1 only examines viability, not ability to compete against other genotypes. Decreased fitness of homozygotes in competition with heterozygotes and wildtypes at some life stage (such as larval competition) could reflect incomplete removal of deleterious mutations during introgression into the CS background prior to carrying out drive experiments since recombination on translocation-bearing chromosomes in Drosophila is reduced throughout the involved arms (67, 68). Alternatively, it could also reflect the acquisition of genetic modifiers during the post-introgression crosses of the translocation stocks required to generate large numbers of homozygotes for population cage experiments. Such modifiers would, in this model, increase the fitness of homozygous carriers in competition with non-carrier homozygotes, but would result in a cost to carriers when in competition with heterozygotes and wildtypes. In either of these models it is unclear why fitness of translocations becomes greater than that of wildtype in later generations. Understanding the basis for these dynamics, and whether they are specific to these translocations, will require further study in other genetic backgrounds, and with other engineered translocations.

## Discussion

Here we report the creation of engineered reciprocal translocations able to drive high threshold population replacement in Drosophila. The tools we used to create translocations in Drosophila ‐ transgene cassettes located on two different chromosomes, a dominant marker created through the act of translocation, a site-specific nuclease able to bring about breakage within each cassette, and unique sequences that can mediate recombination between the two chromosomes ‐ should be portable to other species. This, coupled with the common genetic behavior of reciprocal translocations in diverse species (semisterility in heterozygotes), suggests that translocation-based, high threshold and reversible drive may be possible in many species.

An important unknown from previous work is whether engineered translocations with high fitness are rare or common. Our observations demonstrating population replacement at high but not low introduction frequencies, while limited to two translocations sharing one breakpoint in common, suggest that engineered translocations with high fitness may at least not be rare. That said, while the translocations we generated are competitive in laboratory populations, it remains to be shown that these or any other engineered translocations are fit in competition with the diversity of genotypes that will be encountered in complex natural environments.

Our modeling results suggest that given high enough introduction frequencies, even translocations with high fitness costs, and facing significant levels of incoming migration of wildtypes, can spread to high frequency within a target area. However, modeling also identifies a set of tradeoffs associated with high threshold gene drive. Population replacement is local, but gene flow due to migration has significant effects on the equilibrium frequencies of transgenes within and outside the target area. Consideration of these effects will be important in identifying contexts in which population replacement is likely to have an epidemiological impact, and is able to satisfy regulatory requirements relating to the presence and movement of transgene-bearing organisms. These points on gene flow within a target species notwithstanding, translocation-based drive should be very species specific. This is because drive involves the behavior of entire recombinant chromosomes. It seems unlikely that such a novel entity would thrive when transferred to a different species through mating or horizontal gene transfer.

A key feature of any population replacement mechanism is its degree of evolutionary stability. A translocation drives because its presence in a single copy in heterozygotes creates a toxic condition (genomic imbalance in some gametes) that can be prevented by a second copy of the translocation, which results in the creation of a fit translocation homozygote (genomic balance in all gametes). One can think of this as a toxin-antidote system in which the toxin (the translocation) is dominant (one copy results in genomic imbalance and some death) and the antidote is recessive (two copies of the translocation results in genomic balance and progeny viability). However, in contrast to other toxin-antidote gene drive systems (23, 32, 33, 35-40), the toxin and antidote functions of a translocation are inextricably linked: the toxin is the translocation (in one copy), and the antidote is also the translocation (in two copies). It is presumably very unlikely that the translocation will revert back to the wildtype chromosome configuration. However, even if this happened, necessarily in a single rare individual, this chromosome would be eliminated along with other wildtype chromosomes in a population (of this or any other species (see above)) in which the translocation was present at high frequency. In short, translocation-dependent gene drive cannot break down through mutation of toxin function to inactivity, as with many other chromosomally based drive mechanisms. It is also insensitive to chromosomal sequence variation, mutation and non-homologous end joining, which can prevent the spread of homing-based gene drive mechanisms that rely on cleavage of a specific target sequence (69, 70). Finally, the genes of interest will be placed at the translocation breakpoints. Meiotic recombination is inhibited in these regions (67, 68). In addition, the transgenes are not located in regions that undergo pairing during meiosis. Since they are insertions of novel sequences, they are adjacent to regions that undergo pairing. Thus, transgenes are unlikely to become unlinked from the translocation breakpoint.

Finally, with any population replacement strategy one must plan for the eventual failure of the cargo, whether it encodes one or more genes that mediate disease resistance, or conditional lethality. Failure can occur through evolution of the pathogen. It can also occur through mutational inactivation of the cargo genes. In this latter case, if loss of cargo gene function also results in loss of an associated fitness cost, chromosomes carrying the mutant allele will spread at the expense of those carrying the functional allele. While mutation to inactivity cannot be prevented, chromosome-based drive mechanisms such as translocations have the attractive feature that it should be possible to incorporate multiple transgenes near the breakpoints, bringing about redundancy in effector function and thereby increased functional lifetime in the wild. Cycles of population replacement to bring new genes into the population can also be imagined. In one approach, the translocation can first be removed from the population by driving its frequency below the threshold needed for drive, through dilution with wildtypes. This can then be followed by a second release of a new translocation-bearing strain that has the same breakpoints, and a new cargo. Alternatively, if high fitness translocations with distinct breakpoints can be generated routinely, it may be possible to drive a first generation translocation and any remaining widltypes out of the population in favor of a second, distinct translocation (a point also made by Serebrovskii (47) in the context of use of translocations for population suppression) carrying a new cargo, as with proposals for cycles of replacement of *Medea*-based gene drive systems (5, 21).

The above positive points notwithstanding, several unknowns remain to the implementation of translocation-based population replacement in other insects. First, generating translocations with the approaches described herein will be more challenging in other species in which a high quality annotated genome sequence is not available. Such a resource allows one to identify gene deserts, good candidates for sites in which to locate breakpoints associated with a minimal fitness cost to carriers. It also allows one to determine the orientation with respect to the centromere of sequences that mediate homologous recombination at breakpoints, so as to promote the formation of translocations rather than dicentric and acentric chromosomes. As an example, while the level of annotation of the *Aedes aegypti* genome sequence and transcriptome is otherwise quite high, much of the genome is annotated as a series of contigs of unknown orientation, due to the large amount of repetitive sequences in the genome. Finally, a sequenced genome makes it possible to identify or create, using HEGs, Zinc fingers, TALENs or Crispr/Cas9, site-specific nucleases that promote recombination by cleaving within the transgenes but not elsewhere in the genome.

In addition, the models we have used to characterize translocation behavior do not take into account important real world variables such as non-random mating and local spatial heterogeneity, which can affect the dynamics of translocation spread (55, 71). In order to understand how these and other environmental variables effect translocation-based replacement, and high threshold replacement more generally, it will be important to model drive element behavior using spatially explicit models based on analysis of real populations in complex environments (72, 73). Finally, mosquito populations in the wild consist of multiple chromosomal forms, and may also display some level of reproductive isolation (74-76). How engineered translocations will fare in the face of these variants remains to be determined, but can be explored in competition with genetically diverse laboratory strains (77, 78). While an understanding of the above issues is critical for the success of any population-replacement strategy, the problems are not intractable, as evidenced by successes in controlling pest populations using non-transgenic (79) and transgenic inundative population suppression strategies (80, 81).

## Methods

### Construct Assembly

The Gibson enzymatic assembly (EA) cloning method was used for all cloning (82). For both constructs (A and B), translocation allele components were cloned into the multiple cloning site (MCS) of a plasmid (83) containing the *white* gene as a marker and an attB-docking site. For construct A (Figure 1B), the oenocyte-specific *svp* enhancer (65) and Hsp70 basal promoter fragments were amplified from *Drosophila melanogaster* genomic DNA using primers P16 and P17 (*svp*) and P18 and P19 (Hsp70). The GFP fragment was amplified from template pAAV-GFP (addgene plasmid #32395) using primers P26 and P27. A Kozak sequence (CAACAAA) directly 5’ of the GFP start codon was added with primer P26. The SV40 3’UTR fragment was amplified from template pMos-3xP3-DsRed-attp (addgene plasmid #52904) using primers P28 and P10. The 5’ and 3’ CTCF insulator fragments (84) were amplified from *Drosophila melanogaster* genomic DNA using primers P11 and P15 (for the 5’ CTCF fragment) and P13 and P14 (for the 3’ CTCF fragment). The 667 XYZ and 668 UVW homology fragments were amplified as above with primers P22 and P23 (XYZ) and P20 and P21 (UVW), from plasmid pFUSE-mIgG1-Fc Invivogen, San Diego). The 5’ and 3’ splice sites utilized were from a 67bp intron located in the *Drosophila melanogaster* Myosin Heavy Chain (Mhc) gene ID CG17927. They were added to UVW and XYZ sequences using PCR; the 5’ splice site was added to the 5’ end of the UVW fragment via PCR with primer P24, and the 3’ splice site was added to the 3’ end of fragment XYZ via PCR with primer P25. Two I-SceI recognition sequences Two 18bp I-SceI recognition sequences (ATTACCCTGTTATCCCTA-CTAG-TAGGGATAACAGGGTAAT) were added to the 3’ end of the UVW fragment with primer P21 and the 5’ end of the XYZ fragment with primer P22. The construct was assembled in two steps, as above, with the first (5’) CTCF, the svp and hsp70 fragments, the UVW fragment, and the XYZ fragment cloned in via a first EA cloning step, and the GFP fragment, the SV40 3’UTR fragment, and the second (3’) CTCF cloned in via a second EA cloning step. For construct B (Figure 1B), the opie2 promoter fragment was amplified from plasmid pIZ/V5-His/CAT (Invitrogen) using primers P1 and P2. The XYZ and UVW homology fragments were amplified from plasmid pFUSEss-CHIg-mG1 using primers P3 and P4 (XYZ) and P5 and P6 (UVW). Two 18bp I-SceI recognition sequences (ATTACCCTGTTATCCCTA-CTAG-TAGGGATAACAGGGTAAT) were added to the 3’ end of the XYZ fragment and the 5’ end of the UVW fragment in inverse orientation to each other separated by a 4bp linker sequence (CTAG) using primers P4 (for XYZ) and P5 (for UVW). The 5’ and 3’ splice sites utilized were from a 67bp intron located in the *Drosophila melanogaster* Myosin Heavy Chain (Mhc) gene ID CG17927; the 5’ splice site was added to the 5’ end of the XYZ fragment via PCR with primer P7, and the 3’ splice site was added to the 3’ end of fragment UVW via PCR with primer P8. The dsRed fragment, together with the SV40 3’UTR, were amplified from template pMos-3xP3-DsRed-attp (addgene plasmid #52904) using primers P9 and P10, with a Kozak sequence (CAACAAA) directly 5’ of the DsRed start codon added with primer P9. The 5’ and 3’ CTCF insulator fragments (84) were amplified from *Drosophila melanogaster* genomic DNA using primers P11 and P12 (for the 5’ CTCF fragment) and P13 and P14 (for the 3’ CTCF fragment). The construct was assembled in two steps. First, the *Drosophila melanogaster* attB stock plasmid (83) was digested with AscI and XbaI, and the first (5’) CTCF, the opie-2 promoter, the XYZ fragment, and the UVW fragments were cloned via EA cloning. Then, the resulting plasmid was digested with XhoI, and the dsRed-SV40 3’UTR fragment and the second (3’) CTCF were cloned in via EA cloning. All sequences were analyzed with NNSPLICE 0.9 (available at http://www.fruitfly.org/seq_tools/splice.html) to confirm strength of splice signals and to check for cryptic splice sites. A list of primer sequences used in the above construct assembly can be found in Supplementary Table 1.

### Fly Culture and Strains

Fly husbandry and crosses were performed under standard conditions at 25°C. Rainbow Transgenics (Camarillo, CA) carried out all of the fly injections. Bloomington Stock Center (BSC) fly strains utilized to generate translocations were attP lines 68E (BSC #24485: y^1^ M{vas-int.Dm}ZH-2A w*; M{3xP3-RFP.attP’}ZH-68E), 51C (BSC #24482; y[1] M{vas-int.Dm}ZH-2A w[*]; M{3xP3-RFP.attP’}ZH-51C), and 70A2 (BSC #9741: y[1] w[1118]; PBac{y[+]-attP-9A}VK00023). Fly Stock BSC#6935 (y[1] w[*]; P{ry[+t7.2]=70FLP}23 P{v[+t1.8]=70I-SceI}4A/TM) was used as the source of heat shock induced I-SceI. For balancing chromosomes, fly stocks BSC#39631 (w[*]; wg[Sp-1]/CyO; P{ry[+t7.2]=neoFRT}82B lsn[SS6]/TM6C, Sb[1]) BSC#2555 (CyO/sna[Sco]) were used. For introgression into a wild type background we used the Canton-S stock BSC#1. Translocation construct A was inserted at site 51C, and construct B was inserted at 68E and 70A2 using phiC31 mediated attP/attB integration. These site combinations allowed for the generation of two distinct translocation types, 51C;68E and 51C;70A2. Stocks homozygous for both constructs were then mated with flies that express I-SceI under the control of the Hsp70 heat shock promoter(66). Progeny carrying all three transgenes were subjected to 5 rounds of heat shock during larval stages and as adults. Heat shocks were conducted by submerging fly vials in a water bath set to 38°C for one hour. Adults were outcrossed to w-, and progeny examined under a fluorescent dissecting scope for ubiquitous GFP expression, indicative of translocation generation.

Homozygous translocation-bearing stocks were generated for both 51C;68E and 51C;70A2 site combinations by crossing translocation heterozygotes and identifying homozygous progeny by eye color (light orange eyes for homozygotes versus yellow for heterozygotes for the 51C;68E site combination; light red eyes for homozygotes versus orange for heterozygotes for the 51C;70A2 site combination. After confirming homozygous viability, translocations were introgressed into a Canton-S genetic background. First, CS females were crossed to translocation-bearing males so as to bring the CS mitochondrial genotype into the translocation background. Subsequently, translocation heterozygote females were outcrossed to CS males for 8 generations. Heterozygous translocation-bearing males and virgin females were then crossed to each other to generate homozygous stocks in the CS background for each site combination. Homozygosity was confirmed by outcrossing. Drive experiments for these stocks were set up against CS as the wildtype stock.

### Embryo and Adult viability determination

For embryo viability counts (Table 1), 2-4 day old adult virgin females were mated with males of the relevant genotypes for 2-3 days in egg collection chambers, supplemented with yeast paste. On the following day, a 3hr egg collection was carried out, after first having cleared old eggs from the females through a pre-collection period on a separate plate for 3hrs. Embryos were isolated into groups and kept on an agar surface at 25oC for 48-72 hrs. The % survival was then determined by counting the number of unhatched embryos. One group of 100-200 embryos per cross was scored in each experiment, and each experiment was carried out in biological triplicate. The results presented are averages from these three experiments. Embryo survival was normalized with respect to the % survival observed in parallel experiments carried out with the Canton-S wild-type strain, which was 93.00% ± 1.82%. For adult fly counts (Table 1), individual flies for each genotype cross were singly mated. For each genotype cross, we set up 10-15 individual fly crosses, and the results presented are averages from all these experiments.

### Population cage experiments

All population cage experiments were carried out at 25oC, 12 hour-12 hour day night cycle, with ambient humidity in 250 ml bottles containing Lewis medium supplemented with live, dry yeast. Starting populations for drive experiments included equal numbers of virgins and males of similar ages, for each genotype. Translocation-bearing homozygotes were introduced at population frequencies of 60%, 70%, and 80% (T_1_/T_1_; T_2_/T_2_) for above threshold drive experiments, and 20%, 30%, and 40% (T_1_/T_1_; T_2_/T_2_) for below threshold drive experiments. CS virgin females and males (+/+; +/+) of similar age as the translocation-bearing individuals made up the remainder of the population. The total number of flies for each starting population was 100. All experiments were conducted in triplicate. After being placed together, adult flies were removed after seven days. After another seven days, progeny were collected and divided arbitrarily into two equally sized groups. For one group the fraction of translocation-bearing individuals (T_1_/T_1_; T_2_/T_2_ or T_1_/+; T_2_/+) was determined, while the other group was placed into a new bottle to initiate the next generation.

### Theoretical Framework

We apply the model of Curtis and Robinson (1971) to describe the spread of reciprocal translocations through a population. This is a discrete-generation, deterministic population frequency model assuming random mating and an infinite population size. We denote the first chromosome with a translocated segment by “*T*” and the wild-type version of this chromosome by “*t*.” Similarly, we denote the second chromosome with a translocated segment by “*R*” and the wild-type version of this chromosome by “*r*.” As a two-locus system, there are nine possible genotypes; however, only individuals carrying the full chromosome complement are viable, which corresponds to the genotypes *TTRR*, *TtRr* and *ttrr*, the proportion of the *k*th generation of which are denoted by 
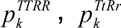
 and 
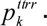
. The four haplotypes that determine the genotype frequencies in the next generation – *TR*, *tR*, *Tr* and *tr* – are described by the following frequencies:

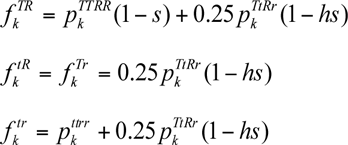

Here, *s* denotes the reduced fecundity of *TTRR* individuals and *hs* denotes the reduced fecundity of *TtRr* individuals relative to wild-type individuals, where *h* ∊[0,1]. By considering all possible mating pairs, the genotype frequencies in the next generation are:

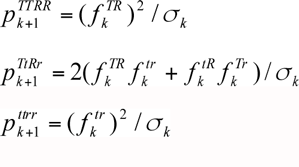

where σ_*k*_ is a normalizing term given by,

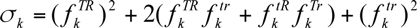

For our three-population models, there are three sets of the above equations to represent each population. We let *m* represent the migration rate per generation. After genotype frequencies for all three populations are calculated for a given generation, a proportion *m* is removed from each genotype from populations 1 and 3 and added to population 2, and a proportion 2*m* is removed from each genotype from population 2, half of which is added to population 1 and the other half of which is added to population 3.

We investigated a number of different fitness cost models and chose the one that provided the best fit to the data. In all cases, the parents in the first generation were not subject to a fitness cost. The simplest model is one in which the fitness of each genotype stays constant over time. Another model considers fitness costs that depend on the population frequency of the genotype. For linear frequency-dependence, this is given by,

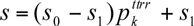

Here, *s*0 represents the fitness cost of a translocation homozygote in an almost fully wild-type population, and *s*1 represents the fitness cost in an almost fully transgenic population. An alternative model is that fitness is time-dependent, as could be explained by introgression of introduced genotypes. For linear time-dependence, this is given by,

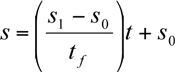

Here, *s*0 represents the fitness cost in the second generation and *s*1 represents the fitness cost in the final generation, denoted by *tf*. For sigmoidal time-dependence, it is given by,

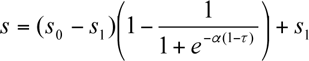

Here, *s*0 and *s*1 are as before, *t* denotes the time of intermediate fitness cost, and *α* denotes the speed of transition between the two fitness costs.

And for exponential time-dependence, it is given by,

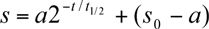

Here, *s*_0_ represents the fitness cost in the second generation, *s*1 represents the fitness cost after many generations, *t*1/2 denotes the time at which the fitness cost is halfway between the two, and *a* is given by,

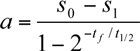

We estimated fitness parameters for each model and compared models according to their Akaike Information Criterion (AIC) values. Model fitting was performed using population count data for the 18 drive experiments conducted for each translocation system (three for each of the 80%, 70%, 60%, 40%, 30% and 20% release frequencies). AIC was calculated as 2*k* – 2log*L*, where *k* denotes the number of model parameters, and the preferred model is the one with the smallest AIC value. The likelihood of the data was calculated, given fitness costs *s* and *hs*, assuming a binomial distribution of the two phenotypes (individuals homozygous or heterozygous for the translocation were considered as the same phenotype to match the experimental counts). Model predictions were used to generate expected genotype proportions over time for each fitness cost, and the log likelihood had the form,

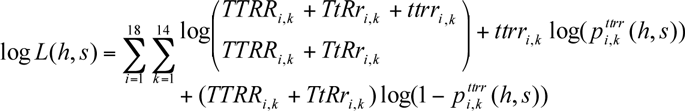

Here, *TTRRi,k*, *TtRri,k* and *ttrri,k* represent the number of *TTRR*, *TtRr* and *ttrr* individuals at generation *k* in experiment *i*, and the corresponding expected genotype frequencies are fitness cost-dependent. The best estimate of the fitness cost is that having the highest log-likelihood. A 95% credible interval was estimated using a Markov Chain Monte Carlo sampling procedure. Matlab and R code implementing these equations is available upon request. The AIC values for each of the fitness cost models are shown in the table below:

**Table 1.**

| **Fitness cost model:** | **AIC (Translocation system 1):** | **AIC (Translocation system 2):** |
| --- | --- | --- |
| Constant fitness costs | 6577.6 | 7448.0 |
| Linear, frequency-dependent fitness costs | 5051.3 | 5572.7 |
| Linear, time-dependent fitness costs | 3888.2 | 3752.1 |
| Sigmoidal, time-dependent fitness costs | 3344.2 | 3321.1 |
| Exponential, time-dependent fitness costs | 3336.2 | 3319.1 |

In summary, the best fitting model for the observed population dynamics is one in which the relative fitness of homozygotes having the translocation is time-dependent, with the relative fitness of these individuals increasing over time, at first rapidly and then converging upon some higher value as described by an exponential function (Figure 5).

**Supplementary Table 1.**
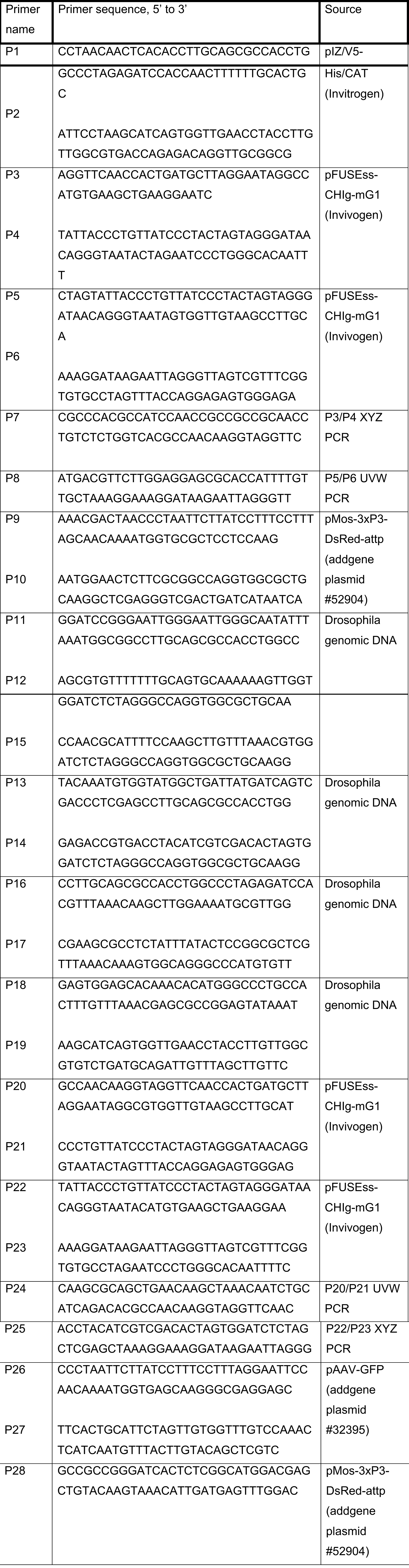
List of primer sequences used in this study.

## Acknowledgements

Work in BAH’s lab (BAH, OSA, ABB, and TI) was supported by the U. S. Army Research Laboratory and the U. S. Army Research Office under contract W911NF-11-2-0055 to the California Institute of Technology, and the USDA and CRDF. Work at UCR (OSA and AB) was supported by an NIH-K22 Career Transition award (5K22AI113060-02) and UCR lab startup funds to OSA. JMM was supported by funding from The Parker Foundation through a gift to the University of California, San Francisco, Global Health Group Malaria Elimination Initiative. TI was supported by NIH training grant 1432.

